# HIV modifies the m^6^A and m^5^C epitranscriptomic landscape of the host cell

**DOI:** 10.1101/2021.01.04.425358

**Authors:** Sara Cristinelli, Paolo Angelino, Andrew Janowczyk, Mauro Delorenzi, Angela Ciuffi

## Abstract

The study of RNA modifications, today known as epitranscriptomics, is of growing interest. The N6-methyladenosine (m^6^A) and 5-methylcytosine (m^5^C) RNA modifications are abundantly present on mRNA molecules, and impact RNA interactions with other proteins or molecules, thereby affecting cellular processes, such as RNA splicing, export, stability and translation. Recently m^6^A and m^5^C marks were found to be present on human immunodeficiency (HIV) transcripts as well and affect viral replication. Therefore, the discovery of RNA methylation provides a new layer of regulation of HIV expression and replication, and thus offers novel array of opportunities to inhibit replication. However, no study has been performed to date to investigate the impact of HIV replication on the transcript methylation level in the infected cell. We used a productive HIV infection model, consisting of the CD4+ SupT1 T cell line infected with a VSV-G pseudotyped HIVeGFP-based vector, to explore the temporal landscape of m^6^A and m^5^C epitranscriptomic marks upon HIV infection, and compare it to mock-treated cells. Cells were collected at 12, 24 and 36h post-infection for mRNA extraction and FACS analysis. M^6^A RNA modifications were investigated by methylated RNA immunoprecipitation followed by high-throughput sequencing (MeRIP-Seq). M^5^C RNA modifications were investigated using a bisulfite conversion approach followed by high-throughput sequencing (BS-Seq).

Our data suggest that HIV Infection impacted the methylation landscape of HIV-infected cells, inducing mostly increased methylation of cellular transcripts upon infection. Indeed, differential methylation (DM) analysis identified 59 m^6^A hypermethylated and only 2 hypomethylated transcripts and 14 m^5^C hypermethylated transcripts and 7 hypomethylated ones. All data and analyses are also freely accessible on an interactive web resource (http://sib-pc17.unil.ch/HIVmain.html). Furthermore, both m^6^A and m^5^C methylations were detected on viral transcripts and viral particle RNA genomes, as previously described, but additional patterns were identified.

This work used differential epitranscriptomic analysis to identify novel players involved in HIV life cycle, thereby providing innovative opportunities for HIV regulation.

## Introduction

The presence of chemical modifications along RNA molecules has been known since the 70s (1). Only recently, however, new technologies allowed the identification and investigation of chemical modifications at transcriptome-wide level, allowing mapping of some modifications in mRNA (2, 3). Similar to epigenetics that focuses on the understanding of DNA and histone modifications in the regulation of transcription, epitranscriptomics investigates RNA modifications and offers a new layer of regulation, impacting and tuning cellular processes, including RNA splicing, export, stability and translation (4). Among these modifications N^6^-methyladenosine (m^6^A) and 5-methylcytosine (m^5^C) are found to be particularly abundant along mRNA molecules (5).

Regulation of RNA modifications is under the control of specific cellular proteins (6, 7). The methylases METTL3-14 together with adaptors proteins act as m^6^A writer complexes of mRNA and catalyze the methylation of adenosine residues within the consensus motif DRA*CH (D = G/A/U, R = G/A, H = U/A/C, and A*= modified A). RNA binding proteins act as m^6^A readers, they bind methylated residues, thereby modulating the fate and metabolism of marked mRNA, *i.e* secondary structure, nuclear export, stability, splicing, and degradation. Demethylases such as ALKBH5 act as erasers of m^6^A, removing the chemical modification from transcripts.

The role and identity of proteins involved in m^5^C turnover is less clear. The addition of m^5^C residues on mRNA molecules is carried out by the methylase NSUN2. M^5^C binding proteins seem to play a role in export and degradation, while to date no m^5^C-specific demethylase has been described yet. The role of RNA modifications is not limited to cellular RNA molecules. Indeed, recent studies highlighted the importance of RNA methylation on viral transcripts as well, including human immunodeficiency virus type 1 (HIV-1, hereafter abbreviated HIV) RNA, and its impact in regulating viral replication and gene expression.

Lichinchi *et al.* reported 14 peaks of m^6^A modification in HIV RNA, including a m^6^A peak in the Rev response-element (RRE) region (8). They showed that RRE methylation increased Rev binding and facilitated nuclear export of viral RNA, thereby enhancing HIV replication. Kennedy *et al.* found four clusters of m^6^A modifications in the 3’ Untranslated region (UTR) of HIV RNA and suggested that the overexpression of the m^6^A readers YTDHF1-3 likely stabilize viral mRNAs, thereby increasing viral replication (9). In contrast, Tirumuru *et al.* and Lu *et al.* showed that HIV RNA has m^6^A modifications in both 5’ and 3’ UTRs, as well as in *gag* and *rev* genes, and that overexpression of YTDHF1-3 proteins in cells inhibits HIV infection by decreasing viral genomic RNA (gRNA) and early reverse transcription products (10, 11).

A recent study from Courtney *et al.* investigate the role of m^5^C in HIV replication (12). Using an antibody-based capture approach, they identified m^5^C-methylated residues in HIV gRNA from CEM T cell-derived virions and on cellular HIV transcripts. They identified the m^5^C mRNA writer NSUN2 as the writer responsible for HIV RNA m^5^C methylation and demonstrated a role of m^5^C in favoring alternative splicing and increasing HIV mRNA translation.

Furthermore, it has been reported that upon HIV infection, the global cellular rate of m^6^A and m^5^C methylation increased (12, 13). However, an in-depth exploration of the differentially methylated genes upon HIV infection is missing.

The discovery of HIV RNA methylation provides a new layer of regulation of HIV expression and replication, and thus a novel array of opportunities to inhibit replication. Investigating the epitranscriptomic landscape of HIV-infected cells will lead to a deeper understanding of HIV-induced RNA modifications and may help to identify novel host cells factors, HIV dependency factors (HDF) or restriction factors (HRF), involved in HIV replication. Indeed, HIV may modulate HDF and HRF to impact viral replication efficiency not only at the level of transcription but also at the level of methylation.

Here we used the SupT1 CD4+ T cellular model infected with a VSV.G pseudotyped HIV-based vector encoding a GFP reporter (HIVeGFP) to explore the m^6^A and m^5^C modification pattern of cellular and viral transcripts in HIV-infected cells, as well as the virion genomic RNA, over time. We found that HIV Infection impacted the methylation landscape of HIV-infected cells by inducing an increased proportion of methylated cellular transcripts. Differential methylation (DM) analysis allowed identifying a few genes that may act as HDF or HRF and thus impact viral replication success. Furthermore, both m^6^A and m^5^C methylation was detected on viral transcripts and on viral particle packaged RNA genome, as previously described, but additional patterns were also detected. All data, at transcriptome, m^6^A and m^5^C epitranscriptome levels, are freely accessible in an interactive mode through HI-TEAM (HIV-Infected cell Transcriptome and EpitrAnscriptoMe), a user-friendly querying platform using an iSee-derived interface (14) available at http://sib-pc17.unil.ch/HIVmain.html.

## Results

### Dynamic analysis of HIV-infected cells

To explore the transcriptomic and epitranscriptomic landscape of HIV-infected cells, we infected SupT1 cells (a CD4+ T-cell-line) with an HIV_GFP-based vector. At 12h, 24h and 36h post-infection, we (i) assessed the percentage of infected cells, monitoring GFP expression by FACS analysis; (ii) measured the amount of viral particles released in the supernatant and (iii) extracted the total RNA, purified polyA RNA and explored the m^6^A and m^5^C landscapes, by either methylated RNA immunoprecipitation sequencing (MeRIP-Seq) or Bisulfite sequencing (BS-Seq) respectively (Figure 1A-B).

**Figure 1.**
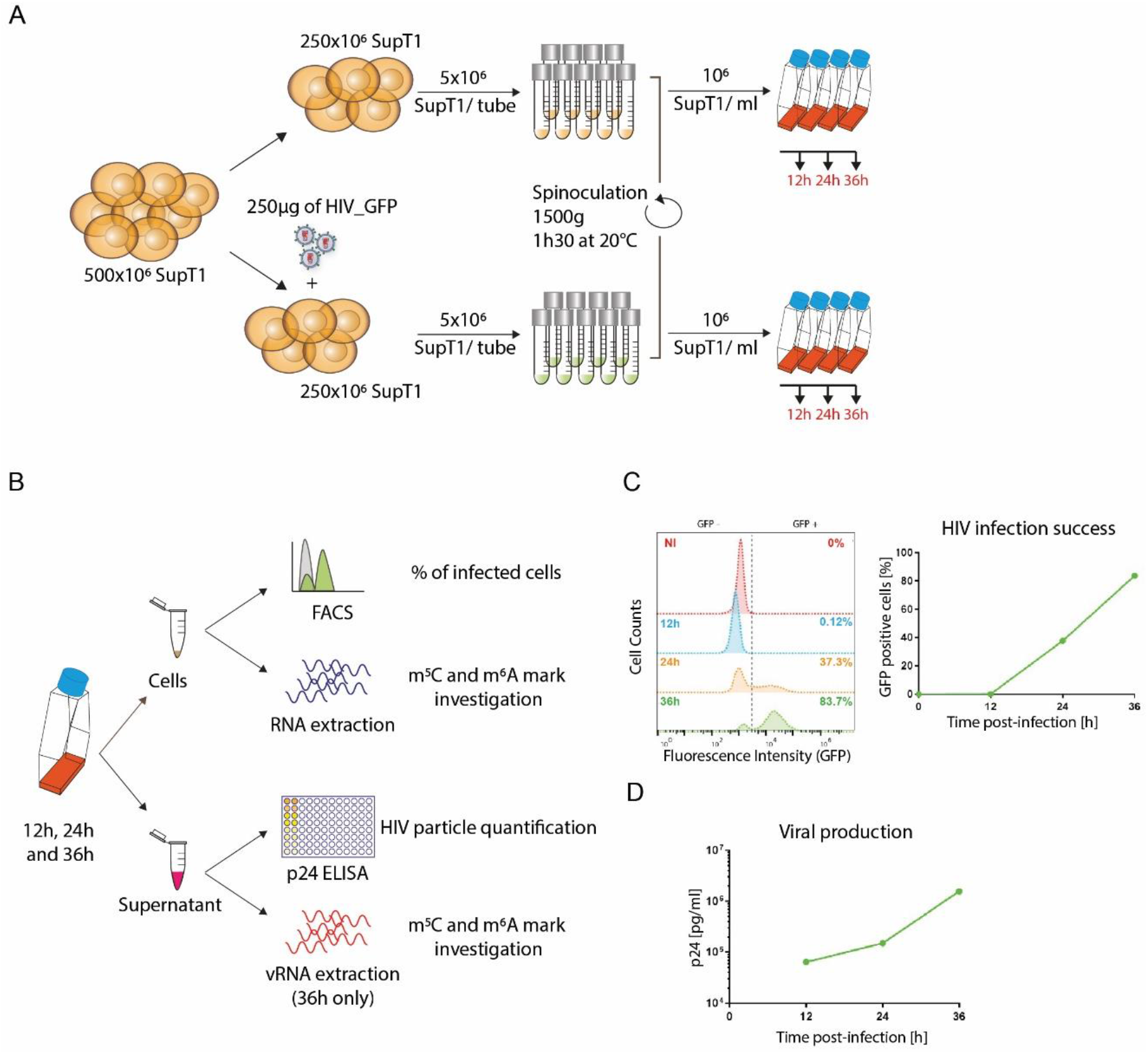
Dynamic analysis of HIV-infected cells. **(A)** Infection setting overview. SupT1 cells were either infected with 1 μg/10^6^ cells p24 equivalent of HIV_GFP virus or left uninfected, divided into aliquots of 5*10^6^ cells/ tube and spinoculated for 90 minutes at 1500 g and 20°C to allow nearly universal infection. Cells were then resuspended at a concentration of 10^6^ cells/ml and further incubated. **(B)** Experimental design overview. At 12, 24 and 36h postinfection, viral supernatant was collected to assess viral production by p24 ELISA; 300.000 cells were fixed and HIV infection success was assessed by evaluating HIV-encoded GFP expression by FACS analysis; the rest of the cells was used for RNA extraction and further analyses on m^6^A and m^5^C epitranscriptomic marks. **(C)** Example of FACS analysis at 12h, 24h and 36h post-infection to evaluate HIV infection success. Left: Histogram plots of FACS analyses showing the GFP intensity (x-axis) on the different samples, non-infected (NI) and infected (12h, 24h, 36h) samples. Right: Graphical representation of the proportion of infected cells (%GFP-positive cells) **(D)** Example of p24 ELISA to monitor viral particle production. Results are expressed as pg of p24 per ml of supernatant over time.

Infection success was monitored over time, following the accumulation of the virally-encoded GFP protein. At 12h post-infection (p.i.), as expected, the GFP expression was not yet detectable, while at 24h p.i. 37.3% of the cells were expressing detectable levels of GFP and 83.7% of the cells were GFP+ at 36h p.i., close to universal infection (Figure 1C). These results were consistent with viral particle production assessed by p24 measurement, which showed increasing viral production over time, with 0.064 x 10^6^ pg/ml at 12 h p.i., 0.150 x 10^6^ pg/ml at 24h p.i. and 1.572 x 10^6^ pg/ml at 36h p.i. (Figure 1D).

### HIV infection induced changes at gene expression level

Transcriptome analysis was performed by RNA-Seq on polyA-selected RNAs over time on infected (HIV) and non-infected mock (NI) SupT1 cells. Altogether, a total of 17,676 genes out of 58,136 were detected (12h NI: 11,908; 12h HIV: 10,980; 24h NI: 13,516; 24h HIV: 12,327; 36h NI: 15,004; 36h HIV: 11,827). In order to increase the specificity of our study we applied a supplemental filter to retain genes above 3 counts per million (CPM). This filter was applied to each condition (Infected or non-infected) individually in order to avoid the introduction of bias upon differential gene expression analyses. Upon quality control and filtering, a total of 13,103 genes was retained for further analysis.

Principal component (PC) analyses separated samples in 2 distinct clusters according to infection condition and time progression, with the PC1 and PC2 representing respectively 67% and 21% of the variance (Figure S1A). Such clustering was not due to the presence of HIV transcripts, as upon their removal, sample distribution was maintained (Figure S1B). Among the 13,103 detected genes, 1,654 (13%) were overexpressed over time while 2,142 (16%) were downregulated. Analysis over time of NI samples only revealed that some genes were differentially expressed, likely due to cell culture conditions and cell growth, but independently from HIV infection. In order to refine the analysis and to observe the *bona fide* impact of HIV infection, the time effect was modeled in a linear model and subtracted to the HIV effect, resulting in improved defined variance (Figure S1C). Thus, upon removal of the time effect, HIV infection alone modulated a total of 1,971 genes, upregulating 813 genes (6.2%) and downregulating 1,158 genes (8.8%) (Figure 2). Gene ontology analysis shows that the 1,971 differentially expressed genes were enriched in the negative regulation of biological and cellular pathways. These data are consistent with our previous study, performed using similar experimental conditions, revealing >75% concordance, and arguing for some degree of reproducibility and confidence (data not shown) (15).

**Figure 2.**
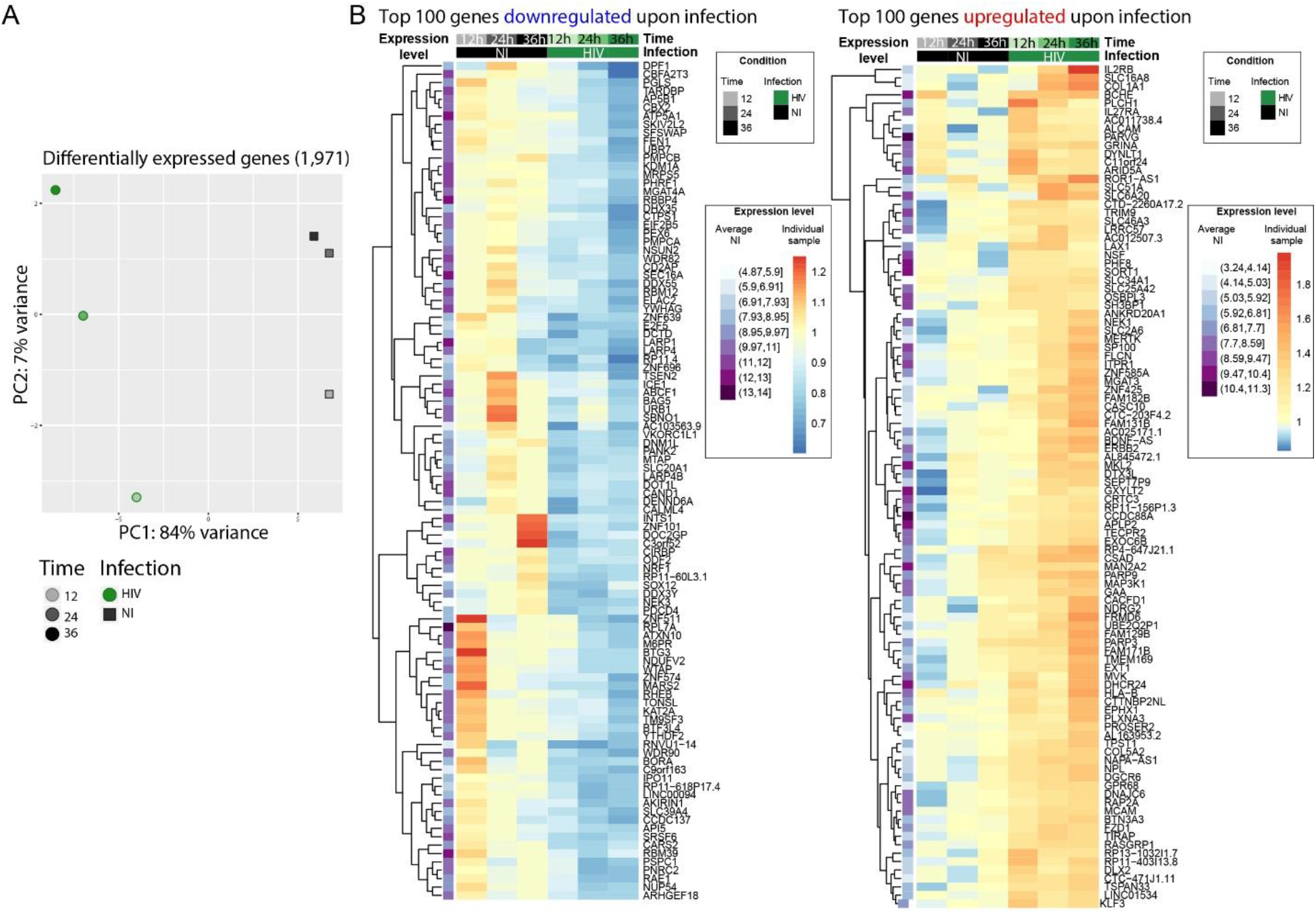
HIV infection induced changes at gene expression level. **(A)** PCA representing 1,971 differentially expressed genes upon HIV infection only. HIV-infected samples (HIV) are represented as green filled circles, non-infected samples (NI) as grey filled squares. Timepoints are depicted by the shade of the color. Time effect has been removed, HIV transcripts are not included. **(B)** Heatmaps of the top 100 (out of 813) downregulated (left) or top 100 (out of 1,158) upregulated (right) genes upon HIV infection. The first column (purple) represents the average gene expression of each gene in the 3 normalized NI samples together, i.e. the darker the color, the higher the expression in the time-averaged NI samples. The log fold change of each gene compared to the average NI gene expression is depicted in shades of blue to red.

Overall, these data confirmed that HIV induced numerous changes at transcriptome level upon infection, that need to be taken into account for an accurate exploration of the epitranscriptomic landscape. Indeed, methylated genes that are strongly impacted by HIV in term of gene expression may introduce a bias to the analysis, *i.e* methylated genes overexpressed upon infection may be considered also as differentially methylated if no correction is applied.

Hence, in order to explore the m^6^A and m^5^C epitranscriptomic landscape of HIV-infected cells independently from their expression level upon infection, all data were normalized to the corresponding gene expression.

### HIV infection induced changes in cellular m^6^A profile

We examined the landscape of the m^6^A RNA methylome during HIV infection at 12h, 24h and 36h postinfection by MeRIP-Seq using either an m^6^A specific antibody or a non-specific IgG antibody as control (16). After pull down and elution, quality and quantity of samples were verified on a fragment analyzer (Figure S2A). The immunoprecipitated RNA was used for library preparation and sequencing; of note, the amount of RNA retrieved from the control condition was too low to perform library preparation and sequencing.

We obtained a range of 26-72 million reads per condition (Figure S2). After quality control and filtering, 8 to 46 million clean reads were kept and further mapped to the HIV and hg38 human genomes, with alignment success typically exceeding 99%.

M^6^A modified regions were identified using the peak calling package MACS2. A total of 17,657 peaks mapping on 7,724 genes across all samples were retrieved representing 59% of the overall detected genes (13,101) (Figure 3A). We looked for the presence of the m^6^A consensus motif DRACH previously identified and detected it in 17,527 peaks out of 17,657 (99.3% of the peaks) (Figure 3B) (5). We further analyzed the 17,527 m^6^A peaks to identify, independently, additional consensus sequences for m^6^A methylation. For this, 20 nucleotides surrounding the center of each m^6^A peak were examined for motif retrieval, and revealed 2 additional highly enriched motifs, WGGAM and GSAGGAGG (Figure S3A); these motif have been previously described as m^6^A binding motif form Zhang et al.(17). As described previously, m^6^A peaks were globally enriched toward the 3’ end of transcripts, and this distribution is not altered upon HIV infection (Figures 3C and S3B) (18). M^6^A modifications were reported to be enriched in long exons (>140 nt), however, upon normalization for exon width, we could not observe significant changes in m^6^A distribution with only a slight enrichment in m^6^A peak frequency in exons >500nt (Figure S3C)(18). Upon PC analysis of the m^6^A peaks retrieved in all samples, we could observe a separation according to time and infection condition, suggesting an impact of HIV infection on the m^6^A methylation profile (Figure 3C).

**Figure 3:**
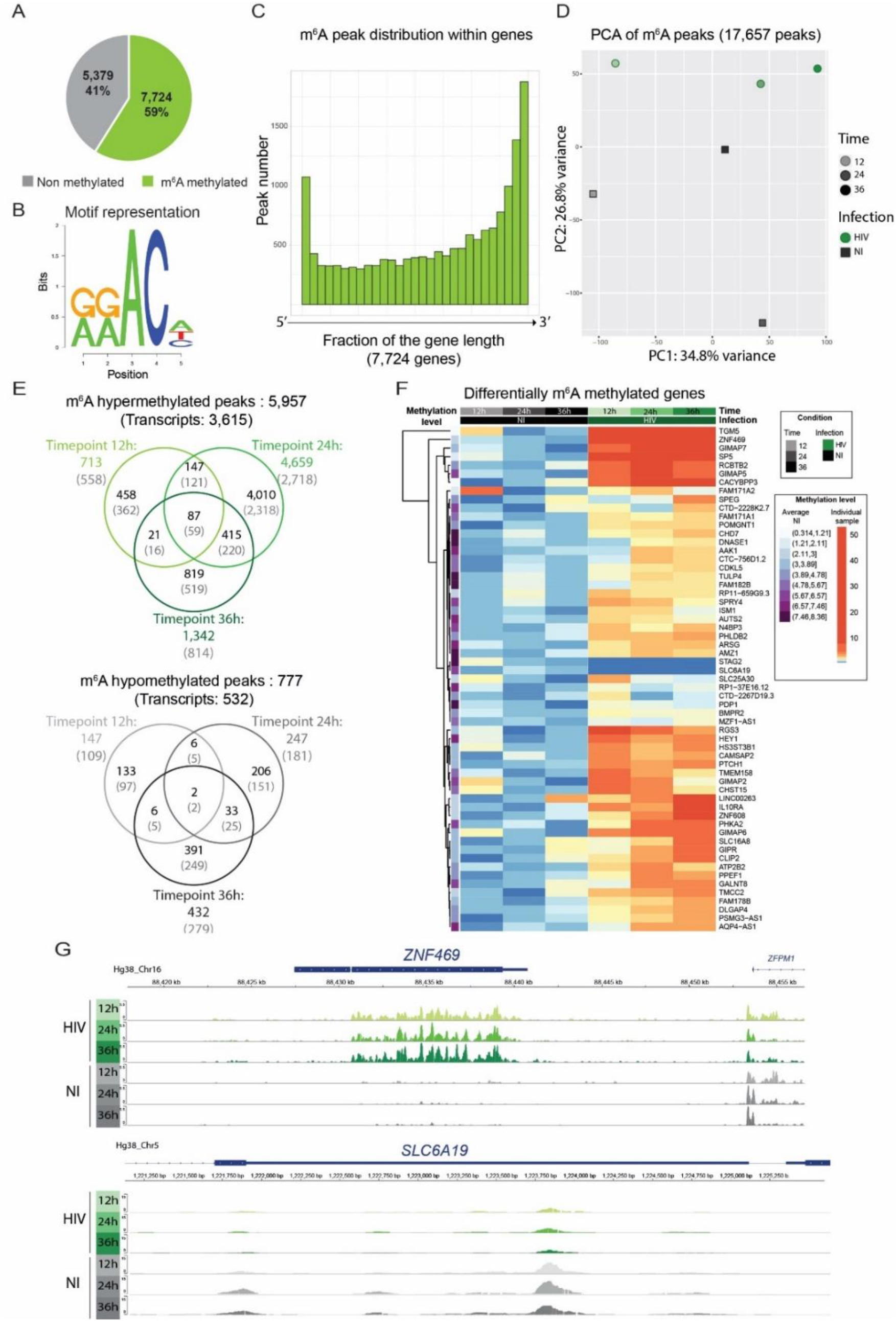
HIV infection induced changes in cellular m^6^A profile. **(A)** Pie-Chart representing the proportion of m^6^A methylated transcripts among the totality of detected transcripts (13,103). **(B)** Representation of the enriched m^6^A DRACH motif among the samples. **(C)** Histogram plots showing on the x-axis genes normalized for their length and divided into 30 bins, and for each bin fraction of the gene, the number of m^6^A residues. **(D)** PCA of the variance of m^6^A peaks among all samples. HIV-infected samples are represented as green filled circles, noninfected samples as grey filled squares. Timepoints are depicted by the shade of the color. HIV transcripts are not included. **(E)** Venn-Diagrams showing hypermethylated (upper panel) or hypomethylated (lower panel) m^6^A peaks upon infection. Values in black represent the number of m^6^A methylated peaks, values in grey into brackets represent the number of corresponding transcripts **(F)** Heatmap of the commonly hyper/hypo methylated transcripts upon infection at the three timepoints. The 61 differentially methylated genes are shown. The average methylation level of the non-infected cells is represented in violet in the first column, and was used for normalization. Differential methylation was then normalized to the average methylation intensity of each transcript. **(G)** Examples of an hypermethylated (upper panel) and an hypomethylated (lower panel) transcript showing m^6^A peak intensity and distribution across samples using IGV viewer.

As m^6^A methylation can occur at different sites along the mRNA molecule, analysis was performed on differentially methylated peaks. A total of 5,957 peaks corresponding to 3,615 transcripts, were found as being hypermethylated upon HIV infection, with 713 peaks at 12h; 4,696 at 24h and 1,342 at 36h post infection (corresponding to 558, 2,718 and 814 transcripts at 12h, 24h and 36h respectively). Only 777 hypomethylated peaks (532 transcripts) were identified, with 147, 247 and 432 peaks at 12h, 24h, 36h post infection, corresponding respectively to 109 transcripts at 12h, 181 at 24h and 279 at 36h post infection (Figure 3D).

We identified 87 m^6^A peaks, representing 59 different transcripts that were commonly hypermethylated in infected cells at 12h, 24h and 36h post-infection. However, only 2 peaks, identified as the stromal antigen 1 (STAG1) and the solute carrier family 6 member 19 (SLC6A19) respectively, were found to be commonly hypomethylated upon infection at the 3 timepoints (Figure 3D to F). Gene ontology analysis of the 61 commonly differentially methylated mRNAs did not reveal any statistically significant enrichment in biological process (data not shown). However, we noticed the presence of 4 out of the 7 GTPase Immuno-Associated Proteins (GIMAP) within the commonly differentially methylated transcripts and overall 6 GIMAPs among the totality of the differentially methylated transcripts (GIMAP1, GIMAP2, GIMAP4, GIMAP5, GIMAP6, GIMAP7). GIMAPs are involved in response to pathogens and have a prominent role in T cell survival and differentiation, consistent with a putative role of these genes on HIV replication (19).

### HIV infection induced changes in cellular m^5^C profile

To obtain a transcriptome-wide landscape of m^5^C profiles, we performed BS-Seq on RNA samples purified from HIV-GFP infected and non-infected SupT1 cells (20). Compared to antibody-based techniques, bisulfite conversion allows higher resolution and higher sensitivity, identifying converted and non-converted cytosines at single nucleotide resolution and providing estimations of the methylation rate of each C residue. To assess efficiency of bisulfite conversion treatment, we used the human 28S rRNA as positive control. Indeed, the C4447 residue of this rRNA is known to be methylated with a frequency of 100%.

Therefore, we spiked-in polyA-depleted RNA in each sample to ensure rRNA representation and presence of 28S rRNA in particular. After bisulfite conversion, a sample aliquot was used for RT-PCR and Sanger sequencing of the C4447 encompassing region of the 28S rRNA (Figure S4A). For all samples we observed a complete C-T conversion along the fragment suggesting the absence of methylation on these C residues, except for the C4447 residue that remains unchanged, confirming the methylation status of this specific C residue (Figure S4B).

After library preparation and high-throughput sequencing we obtained a range of 23-43 million reads/sample (Figure S4C) with a low representation of C and an over-representation of T, consistent with successful unmethylated C-to-T bisulfite conversion (Figure S4D). Reads were processed with the meRan-TK package, specific for RNA bisulfite conversion, taking into account the converted reads to allow genome alignment and mapping (21).

To further assess the conversion rate in each sample, we also included a commercially available pool of non-methylated RNA sequences (ERCC spike-in control) in each sample. ERCC sequence analysis showed an average conversion rate of 99.47%, suggesting that bisulfite treatment was efficient (Figure S4E). Due to the lower quality of bisulfite converted reads with respect to non-converted ones, only transcripts covered with more than 30 reads were retained for further analysis. We could observe different methylation rates among transcripts, hence, to improve the quality of the differential methylation analysis, only cytosines displaying a methylation rate > 20% were used. Overall, we identified 2,267 C residues, corresponding to 947 transcripts, present across all the non-infected timepoints with a methylation rate higher than 20% (7% of overall detected transcripts), 567 m^5^C with a methylation rate higher than 50% and 79 with methylation rate >80% (Figure 4A). To date, no consensus was described for m^5^C methylation. We thus analyzed 10 nucleotides surrounding m^5^C residues displaying a methylation rate greater that 80% and we identified a putative consensus sequence in 500 out of 788 highly methylated m^5^C, representing 63.4% of total hits (Figure 4B).

**Figure 4.**
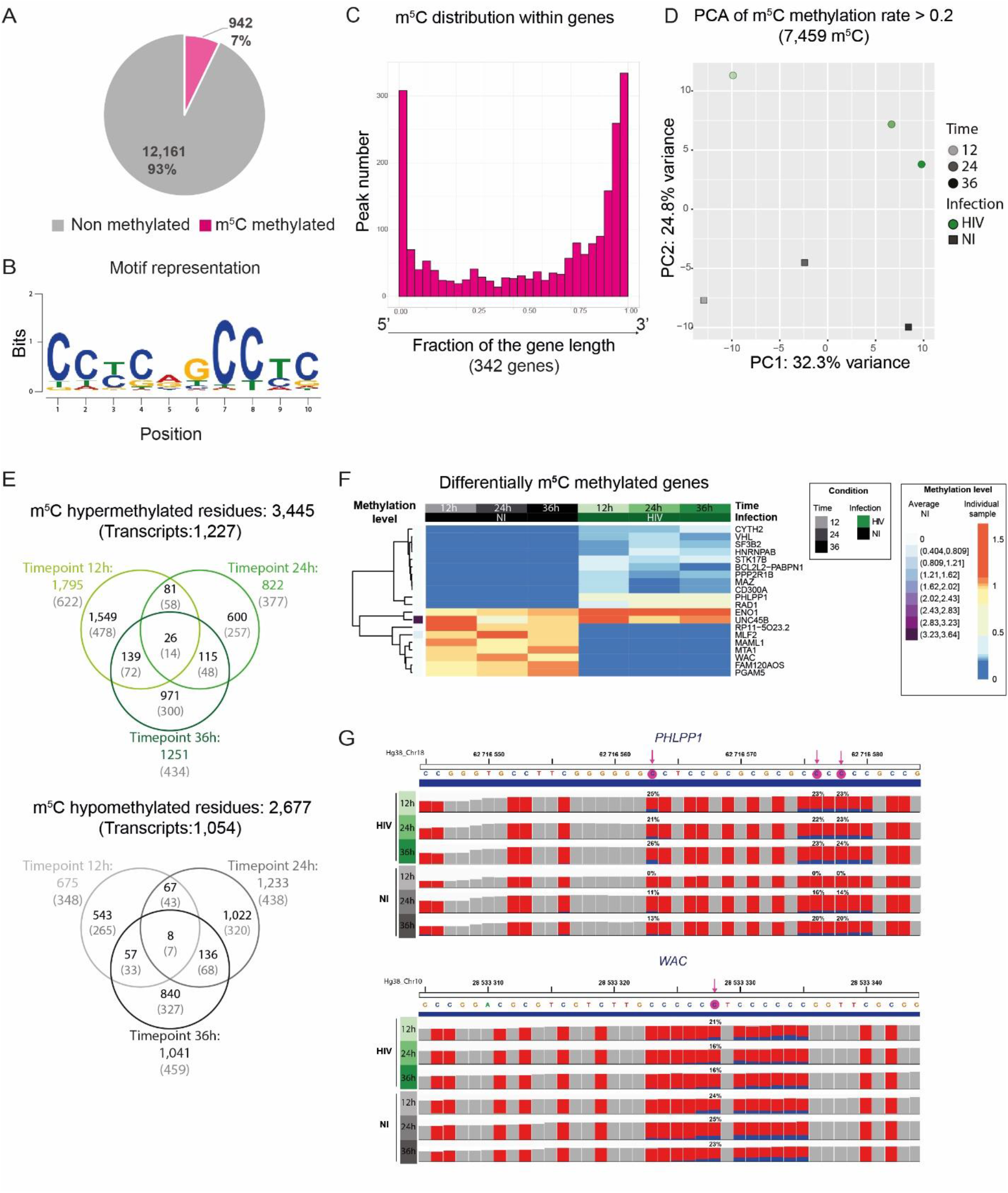
HIV infection induces changes in cellular m^5^C profile. **(A)** Pie-Chart representing the proportion of m^5^C methylated transcripts among the totality of detected transcripts (13,103) **(B)** Identification of a putative consensus motif for C methylation. Logo representation of the predicted m^5^C motif surrounding C residues displaying a methylation rate >80%. **(C)** Histogram plots showing on the x-axis genes normalized for their length and divided into 30 bins, and for each bin fraction of the gene, the number of m^5^C residues. Only genes containing a C-T conversion rate >50% were used. **(D)** PCA of the variance of m^5^C peaks among all samples. HIV-infected samples (HIV) are shown as green filled circles, non-infected samples (NI) as grey filled squares. Timepoint progression is depicted by the shade of the color. HIV transcripts are not included. **(E)** Venn-Diagrams showing hypermethylated (upper panel) or hypomethylated (lower panel) m^5^C residues upon infection. Values in black represent the number of m^5^C residues, values in grey into brackets represent the number of corresponding transcripts. **(F)** Heatmap of the commonly hyper/hypo methylated transcripts upon infection at the three timepoints. The 21 differentially methylated genes are shown. The average methylation level of the non-infected cells is represented in violet on the left, and was used for normalization. Differential methylation was then normalized to the average methylation intensity of each transcript. **(G)** Examples of a m^5^C hypermethylated (upper panel) and a m^5^C hypomethylated (lower panel) transcript upon infection using IGV viewer. Each C residue in the sequence is indicated as a red bar and the proportion of methylated C is shown in blue (exact proportion values are indicated for the statistically significant residues). The significant methylated C residues are highlighted by a pink arrow in the sequence displayed above the tracks.

Consistent with previous studies, m^5^C residues were enriched toward transcript ends and this distribution was not globally affected by HIV infection (Figures 4C and S6) (22).

Principal component analysis performed on the totality of m^5^C shows a separation according to time and infection with 32.3% and 24.8% of the variance explained by PC1 and PC2 respectively. This data remained unchanged upon analysis with a more stringent filter for methylated cytosine proportion (conversion rate >50% and conversion rate >80%) (Figure S7). Altogether, our data suggest that HIV affected the m^5^C profile of cellular transcripts. Upon analysis of differentially methylated m^5^C between infected and non-infected cells, we could identify 1,759 hypermethylated m^5^C in infected cells at 12h; 822 at 24h and 1,251 at 36h post infection (corresponding to 622, 377 and 434 hypermethylated transcripts, respectively). Among them 26 m^5^C mapping on 13 different transcripts (and one unidentified transcript) were commonly hypermethylated in infected cells (Figure 4E). We could also identify 675 m^5^C positions hypomethylated in infected cells at 12h; 1,233 at 24h and 1,041 at 36h post infection (corresponding to 348, 438 and 459 hypomethylated transcripts respectively) with 8 m^5^C mapping on 7 transcripts commonly hypomethylated upon infection (Figure 4E). The hypermethylated and hypomethylated genes common at the three timepoints are displayed in the heatmap (Figure 4E).

Although no statistically significant enrichment was identified by gene ontology analysis, 5 out of the 21 genes (23,8%) identified as differentially methylated were already described as interacting with HIV or contributing to its replication.

### HIV RNA is both m^6^A and m^5^C methylated

Although m^6^A and m^5^C methylation marks were previously reported along HIV RNA molecule, these analyses were performed at a unique time point post-infection and did not consider the putative dynamics of methylation throughout HIV life cycle progression (8–12). We thus took advantage of our temporal design to assess the dynamics of m^6^A and m^5^C epitranscriptomic marks in HIV-infected cells. Furthermore, we compared the methylation profile between intracellular HIV transcripts and vRNA isolated from viral particles.

Upon m^6^A analysis of intracellular HIV RNA molecules, we identified 7 peaks that were conserved at all timepoints, with enrichment of m^6^A peaks toward the 3’ end of the viral sequence (Figure 5A). Increased methylation at the 3’ region was consistent with previous studies identifying the 3’ end as a methylation hotspot and as a binding site for cellular m^6^A readers (9). We also confirmed the presence of two previously reported m^6^A regions in *Pol* (8). However, we identified 2 additional methylated regions, respectively located between *Pol* and *Vif* on one hand, and in *Vpu* on the other hand. Finally, we detected at 36h post-infection a unique peak at the 5’ end of the HIV genome, on the packaging signal sequence psi (ψ), that is also present in the viral particles (Figure S8A).

**Figure 5.**
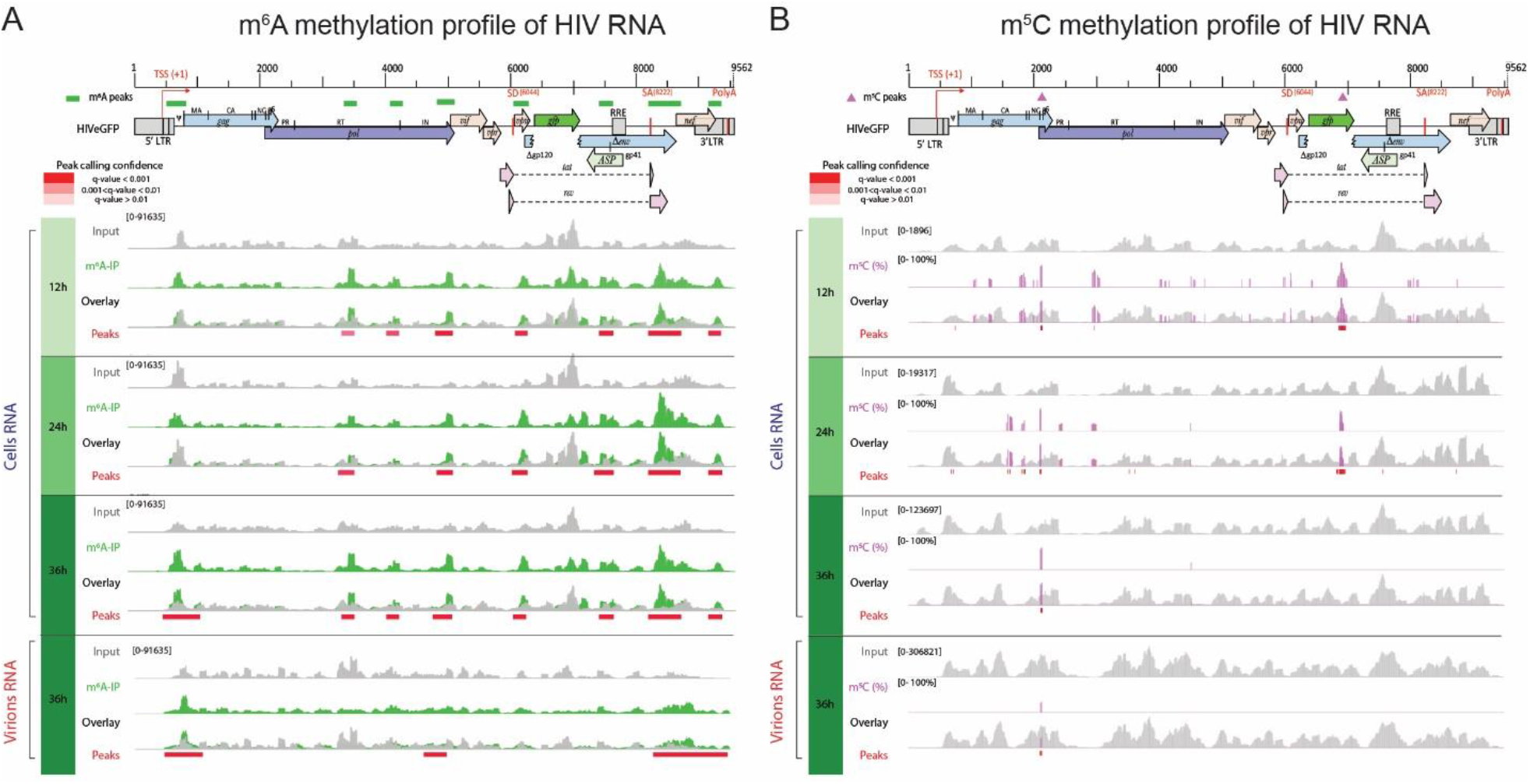
HIV RNA is both m^6^A and m^5^C methylated. Methylation pattern of HIV RNA molecules, isolated from infected cells over time or from viral particles at 36h post-infection. HIV genome organization is depicted on top of the panels and methylation marks are indicated as green color rectangles (A) or pink triangles (B) above the genome, respectively. Detailed read coverage is displayed for each individual sample as tracks below the genome. **(A)** Identification of m^6^A peaks on HIV RNA. Input (gray) and m^6^A immunoprecipitated samples (green) are shown. Putative m^6^A peak calling was performed with MACS2 package after subtraction of the input background (overlay). Statistically significant peaks are highlighted by a red box, with color shading proportional to the q value (m^6^A peak track). **(B)** Identification of m^5^C on HIV RNA. Coverage of HIV genome upon conversion (gray) and detection of m^5^C (pink) are shown. M^5^C are presented as proportion of converted C. Bar height is proportional to the percentage of methylated C in the reads covering the position. The track height is set to 100%. M^5^C calling was performed with MACS2 package. Statistically significant residues are highlighted by a red box, with color shading proportional to the q value.

The methylation pattern found on RNA extracted form viral particles is similar to the one present on intracellular HIV transcripts with the exception of the peak found on *Vpu* in close proximity to the splicing donor site.

Using a bisulfite conversion approach, we confirmed that cellular HIV RNAs were indeed methylated, however with minimal overlap with methylation hotspots described by Courtney *et al.* (Figure 5B) (12). Upon filtering of low coverage regions and statistical analysis we identified 26 m^5^C at 12h post-infection, 30 m^5^C at 24h post-infection and 7 highly methylated m^5^C residues at 36h post-infection covered by at least 100 reads.

Overall, we identified 7 m^5^C residues, common to the 3 timepoints, and clustering all in the vicinity of the HIV *gag-pol* ribosomal frameshift signal with a methylation rate >50%. These highly methylated cytosine are present in viral particles as well as at all timepoints (Figure S8B). The mechanism and the role of this time-dependent effect of m^5^C methylation on the HIV RNA sequence needs further investigation.

## Discussion

Epitranscriptomics is a fast growing field of biology which highlighted the role of m^6^A and m^5^C modifications as specific mRNA marks mostly involved in RNA structural changes and gene expression regulation. In the present study, we explored (i) the cellular m^6^A- and m^5^C-marked transcriptome landscape, (ii) the HIV-induced modifications of the cellular epitranscriptome, and (iii) the position of these specific epitranscriptomic marks on HIV RNA molecule.

Using a SupT1 T cell line infected with a VSV-G pseudotyped HIV-based virus, we detected globally 22.5% transcripts with high confidence (13,103/58,136 genes), among which 15% (1,971/13,103) were differentially expressed, with 813 genes being upregulated and 1,158 downregulated genes in HIV-infected cells compared to mock-treated cells. The analysis of the epitranscriptome, with the tools available today, revealed that 58.9% of genes carried m^6^A methylation (7,724/13,103) (Figure 3B), while m^5^C marks were present on 7% (942/13,103) of transcripts (Figure 4B). These epitranscriptomic marks were mostly enriched towards 3’ ends of transcripts, as shown previously, and this distribution was not affected by HIV infection. Furthermore, our data recapitulated the lower abundance of m^5^C methylation compared to m^6^A modification on mRNA molecules (4). In contrast, in presence of HIV, methylation level globally increased and we identified 62 differentially m^6^A-methylated transcripts (59 hypermethylated and 2 hypomethylated) as well as 21 differentially m^5^C-methylated transcripts (14 hypermethylated and 7 hypomethylated), common at the three analyzed timepoints. Our data are partially consistent with Tirumuru *et al.,* who observed a 4-7 fold-increase of m^6^A methylation in cells infected with a WT virus, but not upon VSV-G-pseudotyped virus infection, suggesting an Env-mediated signaling increase in methylation (13). The basis of this discrepancy is likely due to differences in the experimental design as Tirumuru *et al.* used a global approach, assessing the level of methylation by m^6^A dot-plot on Jurkat T-cell line, while we used the MeRIP-Seq antibody-based technique on SupT1 cells.

Further analysis of the 64 m^6^A-DM transcripts did not reveal any particular enrichment upon gene ontology analysis. Nevertheless, we detected 4 out of 7 GIMAPs in the common list of DM transcripts, and two additional hypermethylated GIMAP members in individual timpoints. GIMAPs are immune-associated proteins displaying a GTPase activity. They have been involved in response to pathogens and have a prominent role in T cell survival and differentiation. The role of GIMAPs in HIV life cycle has never been reported so far and remains to be further characterized.

The analysis of the 21 m^5^C-DM transcripts identified a few genes whose products were previously described as interacting with HIV proteins and affecting the viral life cycle. These include Enolase 1 (ENO1), previously described as hampering HIV reverse transcription (23); the splicing factor 3b subunit 2 (SF3B2), shown to interact with Vpr, thereby impairing splicing of some cellular pre-mRNA and impacting Vpr-mediated G2 cell cycle arrest (24–26); the protein phosphatase 2 scaffold subunit A beta (PPP2R1B), associating with Tat and involved in Tat-mediated apoptosis (27); CD300A, a surface glycoprotein involved in immune response signaling shown to be associated with HIV disease progression markers (28, 29), and shown to be downregulated by Vif (30); and von Hippel Lindau tumor suppressor (VHL), a protein involved in the degradation of hypoxia-inducible-factor and predisposing to cancer when impaired, also known to mediate HIV integrase degradation thereby affecting HIV expression at post-integration steps (31). The role of these methylations on protein expression remains to be investigated, as well as the impact on HIV replication. Nevertheless, these data provide a first roadmap of the impact of HIV on cellular m^5^C-transcriptome.

Altogether, these findings suggest that HIV modulates the host methylation profile of the transcriptome and we can thus hypothesize that the modified transcripts are likely to affect the viral life cycle, either promoting it or inhibiting it. Differentially methylated transcripts may thus represent novel HIV-interacting candidate proteins that should be further investigated and characterized.

Similar to cellular transcripts, the HIV virion RNA molecule is methylated. We identified 7 m^6^A peaks, that were conserved over time, suggesting a rather stable methylation profile. We observed an enrichment of m^6^A at the 3’ of the HIV genome, confirming data from previous studies (8, 9, 11). Our data did not retrieve the two m^6^A peaks previously described to be located in the RRE region, and implicated in enhanced Rev-RRE binding and nuclear export (8). Overall, the studies aiming at investigating m^6^A modifications display minimal overlaps, likely due to protocol differences as mentioned above, and poor resolution of the m^6^A identification approach. Upon comparison between intracellular HIV transcripts and virion RNA we could observe that the m^6^A peak present on *Vpu* and in close proximity of the HIV major 5’ splicing donor (SD) was found only in viral transcripts. Maintenance of the SD hairpin secondary structure is essential to ensure correct splicing of viral transcripts by controlling accessibility of the 5’ splicing site for the splicing machinery (32). Destabilization of the hairpin loop results in an increase of splicing while its stabilization has the opposite effect. We could speculate that the presence of m^6^A in proximity of the site may induce a change in secondary structure allowing easier access to the splicing machinery, while absence of methylation favors the unspliced HIV RNA form. Moreover, the m^6^A peak present in the 3’UTR region in all intracellular viral transcripts is weak or absent in the viral particle genomic RNA, and could suggest a signal contributing to selective packaging of unmethylated HIV RNA genome (Figure S8).

Furthermore, we identified 2 m^6^A peaks, present both on intracellular HIV transcripts and on packaged HIV RNA genome, encompassing the 2 major polypurine tracts (PPT). Although PPT are known for being more resistant to RNAseH-mediated degradation during reverse transcription, the identification of PPT methylation may suggest an additional mechanism providing the observed increased resistance (33). No data were available on m^5^C methylation of HIV transcripts until very recently. Using an immunoprecipitation-based approach to investigate the m^5^C epitranscriptomic mark, Courtney *et al.* identified 18 m^5^C peaks along HIV RNA with an enrichment toward the 3’ end of the genome (12). Using a bisulfite conversion (BS-Seq) approach, we confirmed the presence of this modification on intracellular and packaged genomic viral RNAs and identified 7 conserved, highly methylated m^5^C residues, but with only minimal overlap regarding the exact positions of the epitranscriptomic marks. Using a temporal design, we could describe a C cluster at the beginning of *gag* and surrounding the HIV ribosomal frameshift signal that regulates Gag and Gag-Pol precursor protein synthesis. This signal is indeed essential to maintain a tight regulation of the 20:1 Gag/Gag-Pol translation ratio and ensure successful HIV replication (34). The identification of an m^5^C hotspot close to the frameshift signal may thus point to an additional mechanism involved in the post-transcriptional regulation of Gag and Gag-Pol production. Although m^6^A and m^5^C methylations are considered as the most abundant modifications on mRNA molecules, additional epitransciptomic marks may be present and impact HIV-host interactions, such as 2’-O-methylations (12, 35). Indeed, Ringeard *et al.* recently showed that HIV transcripts can be methylated at the 2’ hydroxyl of ribose, hence 2’-O-methylation, via a specific methyltransferase, FTSJ3, specifically recruited by TAR-RNA binding protein (TRBP) (35). They identified 17 A or U residues containing this specific methylation on the viral RNAs. They demonstrated that 2’-O-methylations were important for viral transcripts to be recognized as endogenous RNA mimics and thus escape innate immune sensing and degradation. Differential analysis of 2’-O-methylation marks upon HIV infection may provide additional insights in HIV life cycle (12, 35).

Overall, this study provided an overview of m^6^A and m^5^C modifications on both viral and cellular transcriptomes over time, identifying the dynamic impact of HIV infection on cellular RNA modifications, and identifying novel candidates as putative factors involved in HIV replication. Further investigation of these candidates, using overexpression or knock-out assays, may reveal a role as HIV dependency factor or inhibitory factor.

The existence of RNA modifications and their potential modulation by HIV proteins offer a new layer of opportunities to hijack the host cellular machinery to promote viral replication and evade the innate immune response. Therefore, identifying all types of differentially methylated or modified transcripts upon HIV infection may lead to the uncovering of novel host factors involved the HIV-host interplay.

## Methods

### Cells and plasmids

Human Embryonic Kidney 293T (HEK293T) cells were cultured in D10 (Dulbecco’s modified Eagle medium (DMEM) containing 1x glutamax (#61965-026, Invitrogen), supplemented with 10% heat-inactivated Fetal Bovine Serum (FBS) and 50 μg/ml Gentamicin) and maintained at a maximal confluence of 80%.

SupT1 cells are human T cell lymphoblasts. They were cultured in R10 (RPMI 1640 with 1x glutamax (#61870-010, Invitrogen) containing 10% heat-inactivated FBS and 25 μg/ml Gentamicin) and split twice a week at 0.5×10^6^ cells/ml to maintain a maximal concentration of 1×10^6^ cells/ml.

The following DNA constructs were used in this study: For viral infection, we used pNL4-3ΔEnv-GFP (NIH AIDS Research and Reference Reagent program, Cat. #11100) that encodes the HIV vector segment with a 903 bp deletion in the *env* ORF in which the *gfp* ORF was introduced. For pseudotyping, the plasmid pMD.G coding for the vesicular stomatitis virus G envelope (VSV-G) was used (36).

### HIV production and infection

For production of HIV-based vector NL4-3-ΔEnv-GFP/VSV-G (named hereafter HIV-eGFP), 2.5 million of HEK293T cells were seeded in 10 cm dishes and incubated over night at 37°C/5% CO_2_. The next day, cells reached about 60 % confluence and were transfected with a total of 10 μg of DNA, *i.e.* 7.5 μg of pNL4-3-ΔEnv-GFP and 2.5 μg pMD2.G coding for the VSV-G envelope, using the jetPRIME kit (Polyplus transfection) and according to manufacturer’s instructions. Briefly, DNA was diluted into 500μl of supplied buffer, mixed with 30μl of jetPRIME reagent and incubated 10 minutes at room temperature. Transfection mixture was then added to the cell dropwise. Fifteen hours after transfection, cells were washed once with D10 and incubated for 33h in 293SFM medium (#11686029, Thermo Fisher Scientific). HIV-GFP particles were harvested 48h after transfection, filtered through 0.45 μm and concentrated on Centricon units (Centricon Plus-70/100K, Millipore). Viral titers were measured by HIV p24 Enzyme-linked immunosorbent assay (ELISA) kit (Innogenetics).

SupT1 cells (5×10^6^ cells) were either mock-treated or infected with 5 μg p24 equivalent of HIV-GFP by spinoculation at 1500 g for 30 min at 20°C, in presence of 4 μg/ml polybrene (Sigma), in 400 μl final volume in 14 ml round bottom polypropylene tubes – a total of 50 tubes were used for mock condition and 50 tubes for infected condition to obtain a total of 250 million cells for each condition. Cells were then pooled, washed three times with culture medium, resuspended at 10^6^ cells/ml in R-10 and further incubated in T75 flasks (8×31ml).

At 12, 24 and 36 hours post-infection, cellular samples (~50×10^6^ cells in 50 ml) were collected for viral and cellular measurements. Briefly, 0.5 ml of the cell cultures were used for cell counting and viability assessment by trypan blue exclusion, using a ViCell Coulter Counter (Beckman Coulter). Remaining cells were centrifuged at 300 g for 10 min. Viral supernatant was collected: 950 μl were mixed with 50 μl NP-40 (0,5%) and stored at −80°C until particle concentration assessment by p24 ELISA (Innogenetics) while the rest of the supernatant was concentrated by filtration on Centricon units (Centricon Plus-70/100K, Millipore) and frozen at −80°C for RNA extraction. Cells were washed with R10 once, centrifuged again, resuspended in 5 ml R10 (~10^7^ cells/ml) and separated as follows: (i) 50 μl of cell suspension were resuspended in Cell Fix 1× (Becton Dickinson) for assessment of GFP expression and infection success by FACS analysis (Accuri C6 FACS, Becton Dickinson), (ii) aliquots of 1 ml of cell suspension (~10^7^ cells) were centrifuged, resuspended in 1ml of Trizol reagent (#15596026, Invitrogen) and stored at −80°C for further RNA extraction and transcriptome analyses.

### RNA extraction

Total RNA was extracted from both concentrated viral particles and cells using Trizol Reagent (#15596026 Invitrogen) according to suppliers’s instructions. Briefly, samples were thawed at room temperature and 200μl chloroform were added to the mixture. Samples were centrifuged for 30min at 10.000g, at 4°C and the RNA–containing, aqueous (upper) phase was transferred to a fresh tube and subjected to precipitation with 0,5 ml of isopropanol for 1h at −80°C. Samples where then centrifuged for 10 min at 12000g, washed once in 1ml of 75% ethanol and resuspended in 50 μl H_2_O.

For poly(A) RNA purification, 200μl Dynabeads Oligo(dT)25 (#61005, Life Technologies) were washed twice with 1ml of binding buffer (20 mM Tris-HCl, pH 7.5, 1.0 M LiCl, 2 mM EDTA) and incubated with aliquots of 75 μg RNA in 100μl final volume for 15 minutes at room temperature on a wheel. Samples were then washed twice with 500μl washing buffer (10 mM Tris-HCl, pH 7.5, 0.15 M LiCl, 1 mM EDTA 10 mM Tris-HCl, pH 7.5), and subjected to a second incubation with the same RNA sample. Poly(A) selected mRNA was recovered through elution by a 2 min incubation with 20μl Tris-HCl (10mM) at 80°C. PolyA depleted RNA from the 36h NI samples was purified and kept as a spike-in control for bisulfite conversion experiments. RNA was purified and concentrated using a column-based kit (#RNA1013, Zymo Research), fragmented during 15 min at 70°C using Ambion RNA Fragmentation Reagents (#AM8740, Life Technologies), in order to obtain fragment of 100-200nt and purified again as above. An aliquot of fragmented RNA (100 ng) was retained as a control for RNA sequencing (input) while the rest was used for MeRIP-Seq and bisulfite conversion allowing m^6^A and m^5^C analysis respectively. At every step, integrity and peak size of the RNA was assessed on a Fragment Analyser (AATI #DNF-472).

### m^6^A-modified RNA immunoprecipitation sequencing (MeRIP-Seq)

For MeRIP (#17-10499, Millipore), 5 μg of fragmented mRNA was incubated with 5 μg of anti-m^6^A antibody or anti-IgG antibody (negative control) previously coupled with 25 μl of A/G-coated magnetic beads in 500 μl IP Buffer for 2h at 4°C following manufacturer’s recommendations. Samples were placed on a magnetic stand for 5 minutes and the unbound RNA was discarded. The beads were then washed three times with 500μl IP buffer and bound RNA was released by two rounds of elution of 1 hour each with 20mM of free m^6^A peptides (7mM N6-Methyladenosine.5’-monophosphate sodium salt). RNA was purified and concentrated in 20μl of water, using a column-based kit (# RNA 1013, Zymo Research). We recovered normally between 15 and 25ng of associated RNA from samples immunoprecipitated with a specific anti-m^6^A antibody. Libraries for sequencing (input RNA-Seq and MeRIP-Seq) were prepared using Illumina TruSeq Stranded mRNA kits (#20020594, Illumina), starting the protocol at the ElutePrime-Fragment step, and with a protocol modification consisting in incubating the samples at 80 °C for 2 minutes to only prime but not further fragment the samples. Samples were sequenced on a HiSeq 2500 Illumina on 4 lanes, using single end reads of 125nt (Genomic Technology Facility (GTF), University of Lausanne).

RNA-Seq data were aligned to a combined hg38 (chr 1-22, X, Y) and HIV genome FASTA using the STAR aligner, and keeping only uniquely mapping reads. Data were analyzed in collaboration with the Swiss Institute of Bioinformatics (SIB) and the Genomic Technology Facility (GTF), University of Lausanne.

### RNA bisulfite conversion sequencing (BS-Seq)

Bisulfite treatment was performed using the EZ RNA methylation Kit (#R5001, Zymo Research). Briefly, 500 ng of poly(A)-selected RNA were spiked-in with 500pg of polyA-depleted RNA (to ensure rRNA representation) as a control for bisulfite conversion. mRNA was mixed with 130μl of RNA conversion solution and converted using three cycles of 10 min denaturation at 70°C followed by 45 min at 64°C in a final volume of 200 μl. After conversion, mRNA was bound to a RNA purification column and desulfonated by addition of 200μl RNA Desulfonation Buffer during 30 minutes at room temperature. Purification was performed using the kit according to manufacturer’s recommendations. RNA quantity and quality was determined by analysis on a Fragment analyser (AATI) using the High sensitivity RNA kit (#DNF-472, AATI).

The efficiency of bisulfite treatment was tested by RT-PCR-mediated bisulfite analysis of spiked-in rRNA (C4447 in 28S rRNA is 100% methylated). Briefly, 4μl of bisulfite converted RNA were subjected to RT with High Capacity cDNA reverse transcription kit (Applied Biosystems #4368814) according to manufacturer procedure and incubated with the following program: 25°C – 10 min; 37°C – 120 min; 85°C 5 min. PCR was performed on 6μl of cDNA using the AccuPrime™ Pfx SuperMix (Thermo Fisher Scientific # 12344-040) with primers annealing on the 28S ribosomal RNA. PCR products were sequenced by Next Generation Sequencing, and resulting sequences aligned to the Human 28S. Cytosine in position 4447 was used as control of non-converted cytosine, while surrounding cytosines were used as a control of CT conversion.

Libraries for sequencing were prepared using the Illumina TruSeq Stranded mRNA kit as described above (*i.e.* entering the protocol at the Elute-Prime-Fragment step and with the modification) and sequenced on two lanes of Illumina HiSeq 2500 as described above.

### FACS analysis

FACS analysis of infected cells was performed on a BD Accuri C6 machine. About 2×10^5^ cells were washed twice in Robosep buffer (#20104, Stemcell Technologies) and fixed in 300 μl CellFix buffer 1X (#340181, BD) for at least 3h at 4°C. The GFP was then monitored by FACS in the FL-1 channel to monitor infection success. Analysis was carried out using FlowJo software.

### Bioinformatic analyses

The analyses described in this section apply to both intracellular transcripts (host mRNAs and vRNAs) and virion-incorporated RNA data.

### m^6^A and gene expression quantification

The m^6^A modification and input libraries underwent a first quality check with FASTQC [http://www.bioinformatics.babraham.ac.uk/projects/fastqc/]. FASTQ files were trimmed with Atropos (37). The following adapter sequences – AGATCGGAAGAG, CTCTTCCGATCT, AACACTCTTTCCCT, AGATCGGAAGAGCG, AGGGAAAGAGTGTT, CGCTCTTCCGATCT – were removed after trimming of low-quality ends (a Phred quality cutoff of 5 has been applied) as specified by the manufacturer (https://support.illumina.com/downloads/illumina-adapter-sequences-document-1000000002694.html). Only reads with a minimum length of 25 base pairs after trimming were retained. Trimmed reads were aligned to an assembly of the Hg38 human genome and HIV [Integrated linear pNL4-3ΔEnv-GFP] genome. The software used for the alignment was HISAT2 (38). Aligned reads were indexed and sorted with SAMtools (39).

Post-alignment quality of the reads was performed with SAMtools stat and Qualimap 2 (40). Quality measures have been collected and summarized with multiQC (41).

HIV genome has homologous 634 bp sequences in the 5’ LTR and 3’ LTR. Multimapping reads from 5’

LTR have been realigned to the corresponding 3’ LTR region with SAMtools.

Abundance quantification of transcripts on input libraries has been performed with Salmon (42). HIV expression level has been quantified by directly counting reads mapping to the viral genome.

m^6^A peaks were identified with the peak calling software MACS2 (v 2.1.2)(43). Caution was applied in the choice of MACS2 running parameters, to allow the toll to correctly work on RNAseq data. In RNA-Seq data the peak calling can be affected by the gene expression level, and short exons may potentially be miscalled as peaks. Hence, signal from input must be subtracted from m^6^A signal, without the the smoothing routinely applied by MACS2 to DNA based data.

‘callpeak’ sub-command from MACS2 was run with the following parameters: –keep-dup auto (controls the MACS2 behavior towards duplicate reads, ‘auto’ allows MACS to calculate the maximum number of reads at the exact same location based on binomial distribution using 1e-5 as p-value cutoff), -g 2.7e9 (size of human genome in bp), -q 0.01 (minimum FDR cutoff to call significant peaks), --nomodel (to bypass building the shifting model, which is tailored for ChIP-Seq experiments), --slocal 0 —llocal 0 (setting these 2 parameters to 0 allows MACS2 to directly subtract, without smoothing, the input reads from the m^6^A reads), --extsize 100 (average length of fragments in bp), -B -SPMR (to generate library size normalized bedGraph track for visualization).

In order to compare infected vs non-infected samples, the differential peak calling sub-command of MACS2, ‘bdgdiff’, was used. ‘bdgdiff’ takes as inputs the bedGraph files generated by ‘callpeak’. First, we run ‘callpeak’ with the same parameters as above, but without the -SPMR option (output unnormalized tracks), which is not compatible with ‘bdgdiff’. Then, for each time point we run the comparison of infected versus non-infected samples with ‘bdgdiff’, subtracting the respective input signal from the m^6^A signal and providing the additional parameters -g 60 −1 120.

### Bisulfite conversion analyses

Cutadapt (44) was applied for read trimming, using parameters of —minimum-length=25 and the adapter “AGATCGGAAGAGCACACGTCTGAAC”. Trimmed reads were subsequently reverse complemented using seqkit (45).

Quality control was performed by employing FastQC to examine samples for (a) poor read quality, and (b) contamination of which there was no supporting evidence.

The application meRanGh from the meRanTK package (21) was leveraged to make an index file for alignment consisting of the hg38 reference genome supplemented with the HIV genome. Aligning again employed meRanGh with parameters enabling unmapped reads (-UN), multi-mapped reads (-MM) to be written to output files. Additionally, the output bedGraph (-bg) was produced.

Reported Regions were filtered by those with at least a 10 read coverage (-mbgc 10). To account for HIV LTR regions being multi-mapped, and not thus not being present in the alignment output file, Sambamba (46) merge was employed to filter reads in the HIV genome upstream of the 8500bp locus and append them to the final alignment.

FeatureCounts (47) was employed at the exon and CDS level for the hg38 and HIV genomes, respectively.

Methylation calling was completed via the meRanCall tool, provided by meRanTK, with a read length (-rl) parameter of 126, an error interval of .1 used for the methylation rate p-value calculation (-ei), an expected conversion rate of .99 (-cr). MeRanCompare was employed with a significance value of .01 as the minimal threshold for reporting. For its size factors parameter, MeRanTK’s included utility estimateSizeFactors.pl was employed on each of the time points, and produced values of (.8102, 1.2342), (1.1894,.8408), (0.6562,1.5240) for (not infected, infected) across time points 12, 24, and 36h respectively.

### Differential Gene Expression (DGE) analysis

Transcript abundance and counts estimated by Salmon for the input samples were imported into an R session (version 3.5.1) using the package tximport (48). The same package was used to summarize transcript level expression at the gene level.

Low count genes have been removed with the method ‘filtered.data’ from the package NOISeq (49). ‘filtered.data’ method 1 removes those genes that have an average expression per condition less than 3

CPM (Counts Per Millions) and a coefficient of variation per condition higher than cv.cutoff = 100 (in percentage) in all the conditions.

The filtered gene table was the processed with the package for differential gene expression analysis (50). First, exploratory PCA plots were generated with the PCA plot function on counts transformed with the rlog method in DESeq2. Then, differential expressed genes were called with an adjusted (Benjamini-Hochberg method) p-value threshold of 0.01. To takeinto account the effect of cell culture time together with that of HIV infection, we asked DESeq2 to fit a Generalize Linear Model (GLM) which included both effects: design = ~ infection + time. Two lists of differentially expressed (DE) genes according to infection and time were thus obtained. To further separate the effect of the HIV infection from the time one, we produced a list of ‘HIV only’ DE genes, by removing from the list of infection related DE genes those in common with the list of time-related DE genes.

A PCA plot with this ‘HIV only’ DE gene list was produced in order to highlight the effect of HIV infection, and heatmaps with the gene expression level of these genes were also drawn.

### m^6^A differential peak calling analysis

MACS2 ‘callpeak’ generated a list of peaks for each time point and each infection status (infected and non-infected). MACS2 ‘‘bdgdiff’ generated 3 lists (common peaks, up and down regulated upon HIV infection) for each time point comparison. These lists of peaks were further processed and analyzed with the R package diffbind (51), and annotation with overlapping genes was provided by the package ChIPpeakAnno (52).

To reduce the number of false positives, only the peaks called by both MACS2 methods (‘callpeak’ and ‘bdgdiff’) were retained in the following analyses. For purpose, for each time point and infection status, we intersected the list produced by ‘callpeak’ with the corresponding lists produced by ‘bdgdiff’ (the common peaks and condition specific peaks). We thus obtained a high confidence peak list for each time point and condition.

We defined a measure for peak intensity based on the number of reads overlapping with each peak. For counting the overlapping reads, the function dba.count from DiffBind was used. First, we created a consensus peak set with the union of the high confidence peak lists. Reads overlapping with a span of 100 bp around the summit of the peaks in the consensus were counted, normalization factors were computed using edgeR TMM method (53), and the reads in the m^6^a input were subtracted to separate methylation from expression level effects. The normalized counts at each peak, which we will call peak scores, were used to generate the PCA plot, the peak distribution along gene length, and heatmaps.

The presence of the m^6^A binding motif (“DRACH”) was assessed using the function scan_sequences from the package “universalmotif” (54) over the consensus list of peaks.

An unsupervised motif search was also performed. From the consensus peak set, we extracted the nucleotide sequence (from the reference genome Hg38) of an interval of 10bp upstream and 10bp downstream from the center of each peak. The list of 17657 sequences was used as input for the tool DREME (55), from MEME suite (5.1.0)(56), which performed the motif discovery.

Peak distributions along genes were computed by dividing each gene in 30 intervals and adding up the scores of peaks belonging to each interval for all genes (in other words, computing the sum of the peaks in each interval weighed by the scores). The distributions were plotted at each time point and condition. In order to compare the modification of m^6^A RNA methylation specific to HIV infection, we intersected the up (down) regulated peak lists of all 3 time points, and, for late infection response, at 24h and 36h time point only. We summarized these results at the gene level (obtaining a ‘gene methylation score’), by adding up the scores of the peaks in each gene. The methylation score of up and down methylated genes upon HIV infection were plotted as heatmaps.

### m^5^C differential methylation calling analysis

The m^5^C data analysis follows the line of the m^6^A one described above. The lists of methylated C generated by meRanCall tool were further processed and analyzed with the R package DiffBind and annotation for overlapping genes was provided by the package ChIPpeakAnno.

In order to reduce the number of false positives in m^5^C called bases, beside the adjusted p-value threshold of 0.01 applied by meRanCall, we introduced an extra threshold on coverage, asking that the retained m^5^C bases having at least 30 read coverage. This number was adjusted by the total number of reads in each library to have an even filter across samples.

Furthermore, a consensus set of m^5^C sites was created by the union of the m^5^C called bases from all samples, asking that a methylated site appears in at least 2 samples. The methylation rate (number of methylated C over total number of C) at each base was used as methylation intensity score to generate the PCA plot, the m^5^C distribution along gene length, and heatmaps.

A motif discovery was performed with MEME (5.1.0)(56). A list of 788 sequences of 10 bp surrounding both sides of methylated bases was input to MEME. This list is a high confidence list of methylated sites made by joining (union) the bases with a methylation rate greater than 0.8 from all samples.

m^5^C distributions along genes were computed by dividing each gene in 30 intervals and adding up the methylation rate of m^5^C belonging to each interval for all genes (in other words, computing the sum of the m^5^C sites in each interval weighed by the methylation rate). The distributions were plotted at each time point and condition.

In order to compare the modification of m^5^C RNA methylation specific to HIV infection, we intersected the up (down) regulated peak lists of all 3 time points, and, for late infection response, at 24h and 36h time point only. We summarized these results at the gene level (obtaining a ‘gene methylation score’), by adding up the methylation rates of the bases in each gene. The methylation scores of up and down methylated genes upon HIV infection were plotted as heatmaps.

## Data availability

All raw RNA-Seq data will be deposited in NCBI’s Gene Expression Omnibus and are accessible through GEO series accession number GSE157193. All data are freely accessible on the iSEE-Hi-TEAM web resource at http://sib-pc17.unil.ch/HIVmain.html.

## Acknowledgements

We thank the Lausanne Genomic Technology Facility of the University of Lausanne. We thank Raquel Martinez for technical assistance and valuable help. This work was supported by the Swiss National Science Foundation (grants 31003A_166412 and 314730_188877).

## Supplementary Figures and Data

### Supplementary Figures

**Figure S1:**
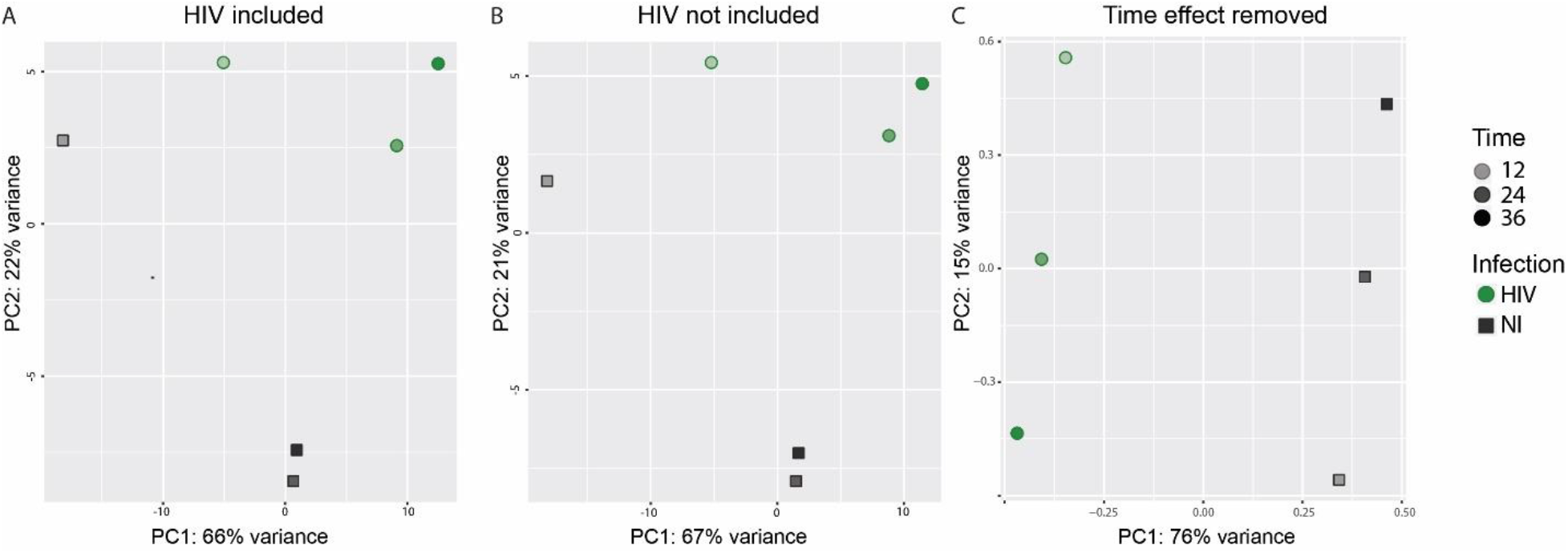
Time and HIV infection effects on gene expression. Principal component analysis (PCA) representing gene expression variation across samples, in presence (A) or absence (B) of HIV transcripts and upon removal of time effect (C). HIV-infected samples (HIV) are represented as green filled circles, non-infected samples (NI) as grey filled squares. Increasing timepoints (12h, 24h, 36h) are depicted by increasing color shading.

**Figure S2.**
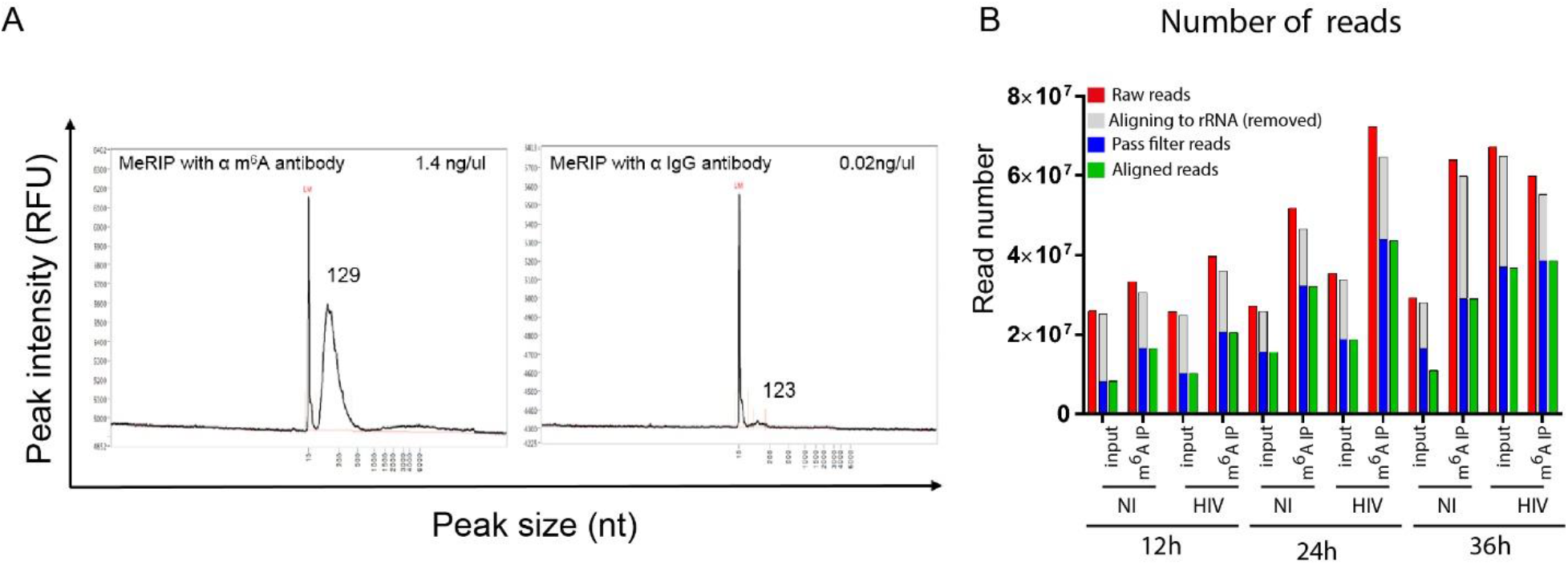
M^6^A quality controls. **(A)** Representative Fragment Analyzer outcome assessing the size and the quantity of RNA fragments recovered after MeRIP-Seq with a specific anti-m^6^A antibody (left) and with the non-specific anti-IgG antibody (right). The x-axis represents the nucleotide size of the peak, while the y-axis quantifies the intensity of the peak in Relative Fluorescence Units (RFU). LM: low marker reference. **(B) Sequencing read analysis.** Amount of raw reads (red), reads aligning to rRNA (grey), clean pass filter reads (blue) and aligned reads (green) retrieved for input and m^6^A-immunoprecipitated (IP) samples. HIV: HIV-infected samples; NI: non-infected mock samples.

**Figure S3.**
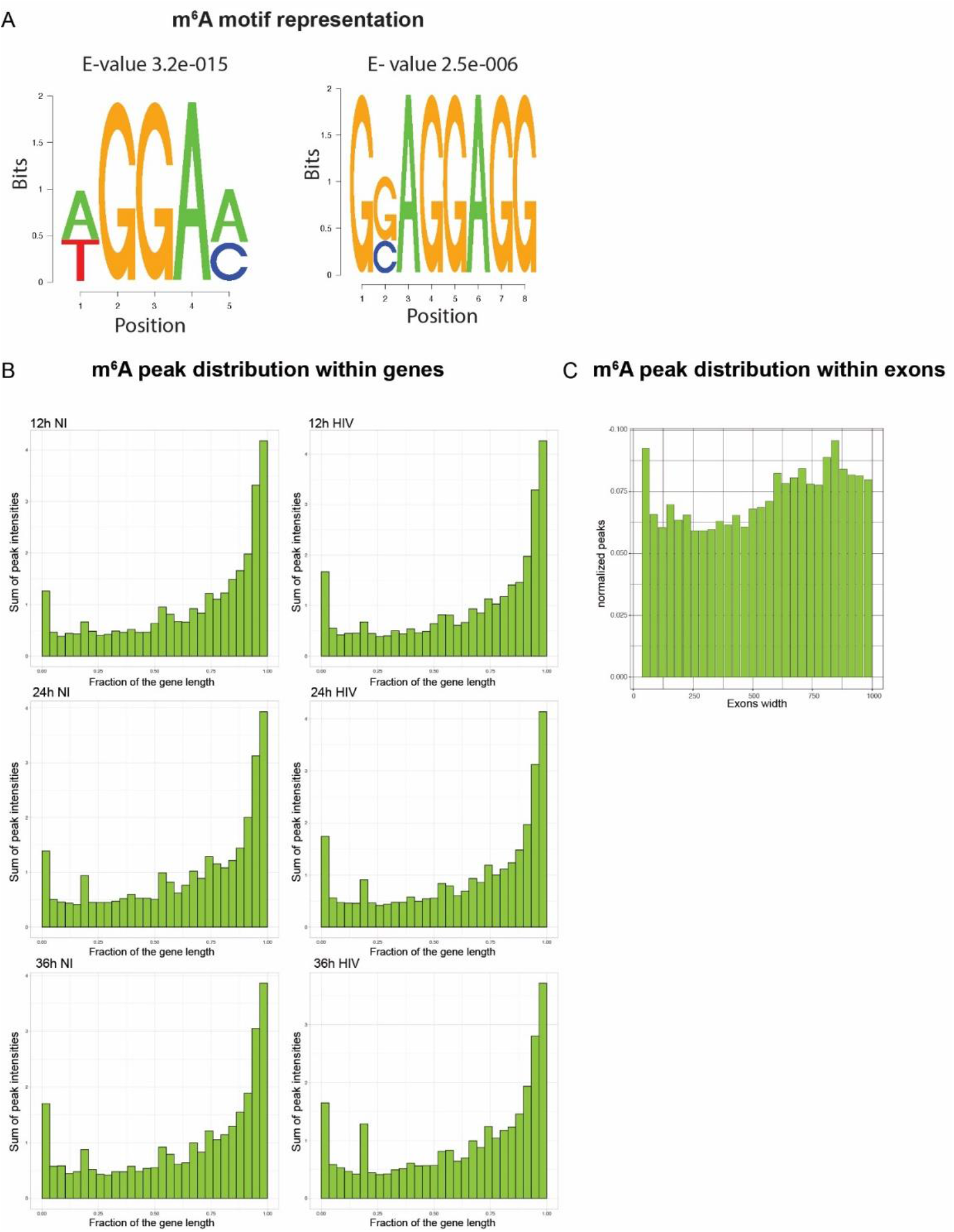
m^6^A peak distribution. **(A)** Representation of the two most enriched m^6^A motif in our dataset **(B)** Frequency of m^6^A methylation along the gene in each sample. Histogram plots showing on the x-axis genes normalized for their length and divided into 30 bins, and for each bin fraction of the gene, the number of m^6^A peaks weighted for the peak intensity (peak surface) was assessed. **(C)** Frequency of m^6^A peak distribution according to exon width. Exons were binned according to their width (x-axis). The number of m^6^A peaks (y-axis) per binned exon width was normalized by the exon width.

**Figure S4.**
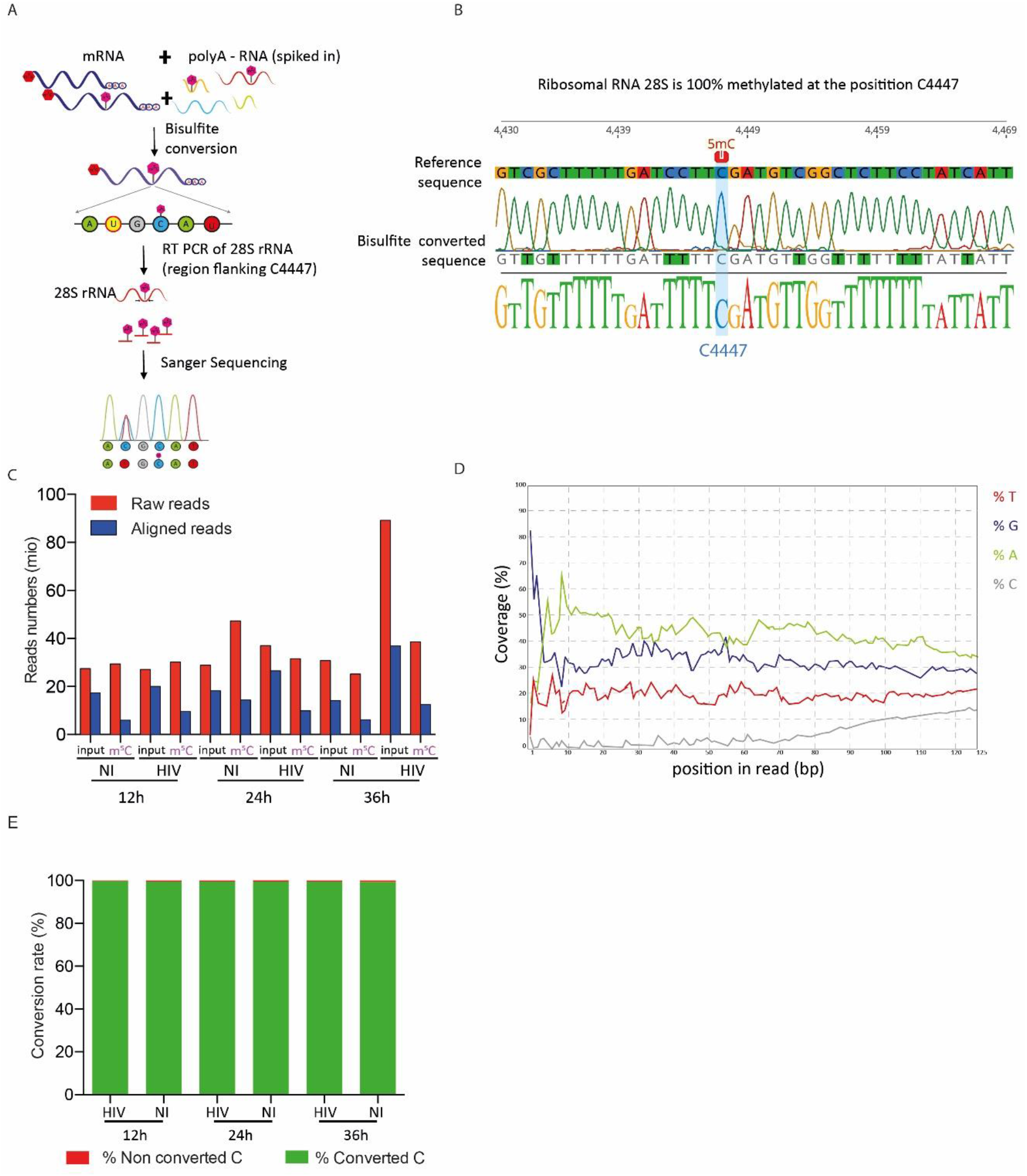
M^5^C quality controls. **(A)** Schematic representation of the evaluation of bisulfite conversion efficiency by qPCR. The 100% methylated C4447 of the 28S human rRNA was used as a positive control to evaluate the efficacy of bisulfite conversion treatment. Purified mRNA was spiked-in with polyA-depleted RNA, to ensure rRNA representation, and exposed to bisulfite treatment, triggering C to U conversion in absence of methylation while methylated C (pink hexagone mark) were protected and unmodified. A fraction of the bisulfite converted RNA was subjected to RT-PCR in order to amplify a 200 bp region of the 28SrRNA flanking C4447 for Sanger sequencing. **(B)** Electropherogram of a representative sequence retrieved after RT-qPCR. The blue arrow highlights the conserved methylated C residue at position 4447, identified as a C upon sequencing (top panel). In contrast, non-methylated C residues (blue in the reference sequence on the bottom panel) are converted to T (green nucleotides, as shown on the top panel) upon bisulfite treatment, RT and sequencing. **(C)** Amount of raw reads (red) and clean pass filter reads (blue) retrieved for the bisulfite converted samples. HIV: HIV-infected samples; NI: non-infected mock samples. **(D)** Representation of the read per base content along every position of the read. On the x-axis is depicted the position in the read, on the y-axis the total coverage proportion of each base. **(E)** Conversion rate assessment. Non-methylated ERCC control sequences were spiked in each sample to assess the conversion rate. Percentage of successfully converted C are represented in green, non-converted C are represented in red. The average conversion rate across samples was 99.47%.

**Figure S5.**
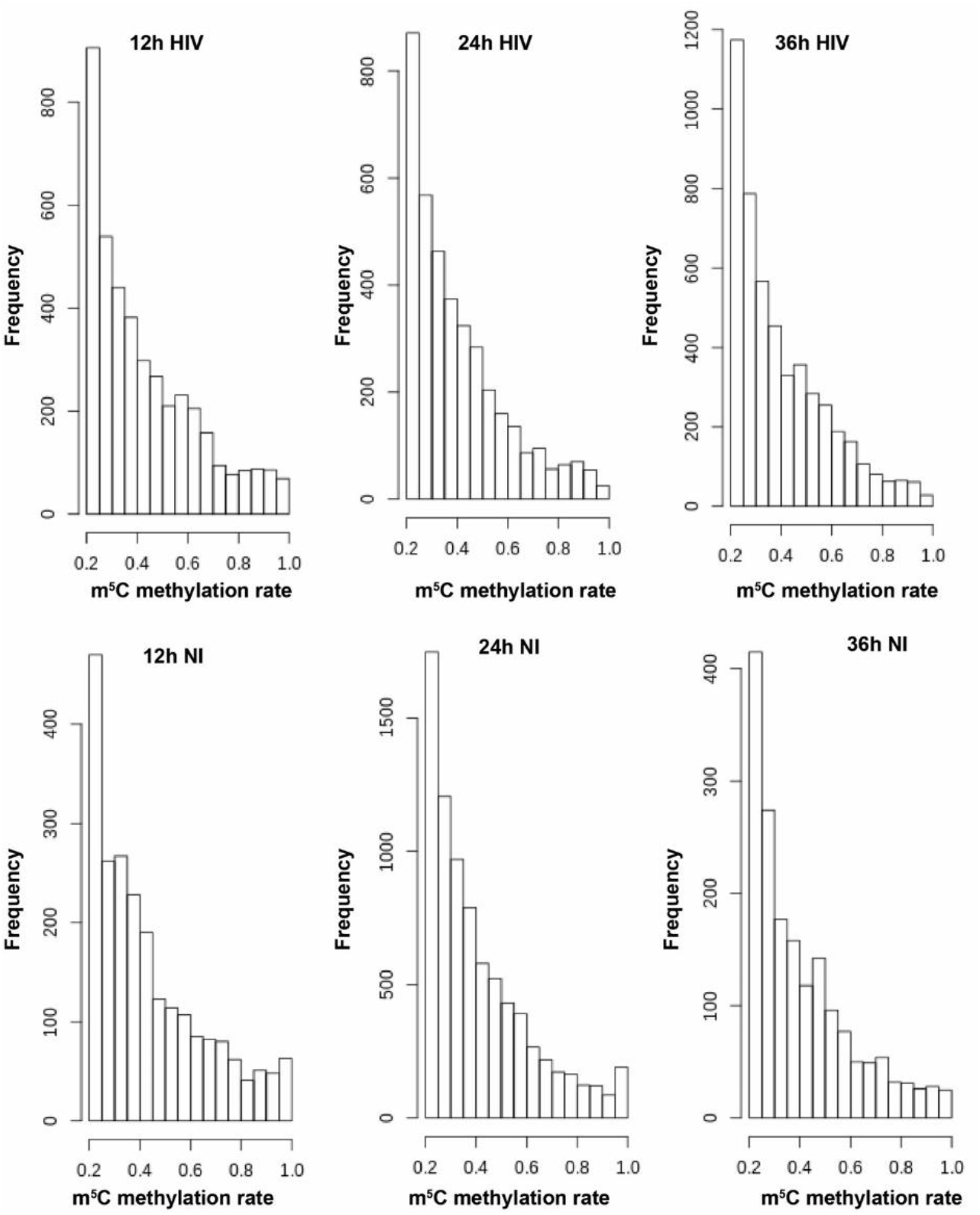
Frequency of m^5^C methylation. Histogram samples showing the number of C residues (y-axis) and their methylation rate (x-axis), *i.e.* the fraction of non-converted (methylated) C, for each sample (infection condition and time condition).

**Figure S6.**
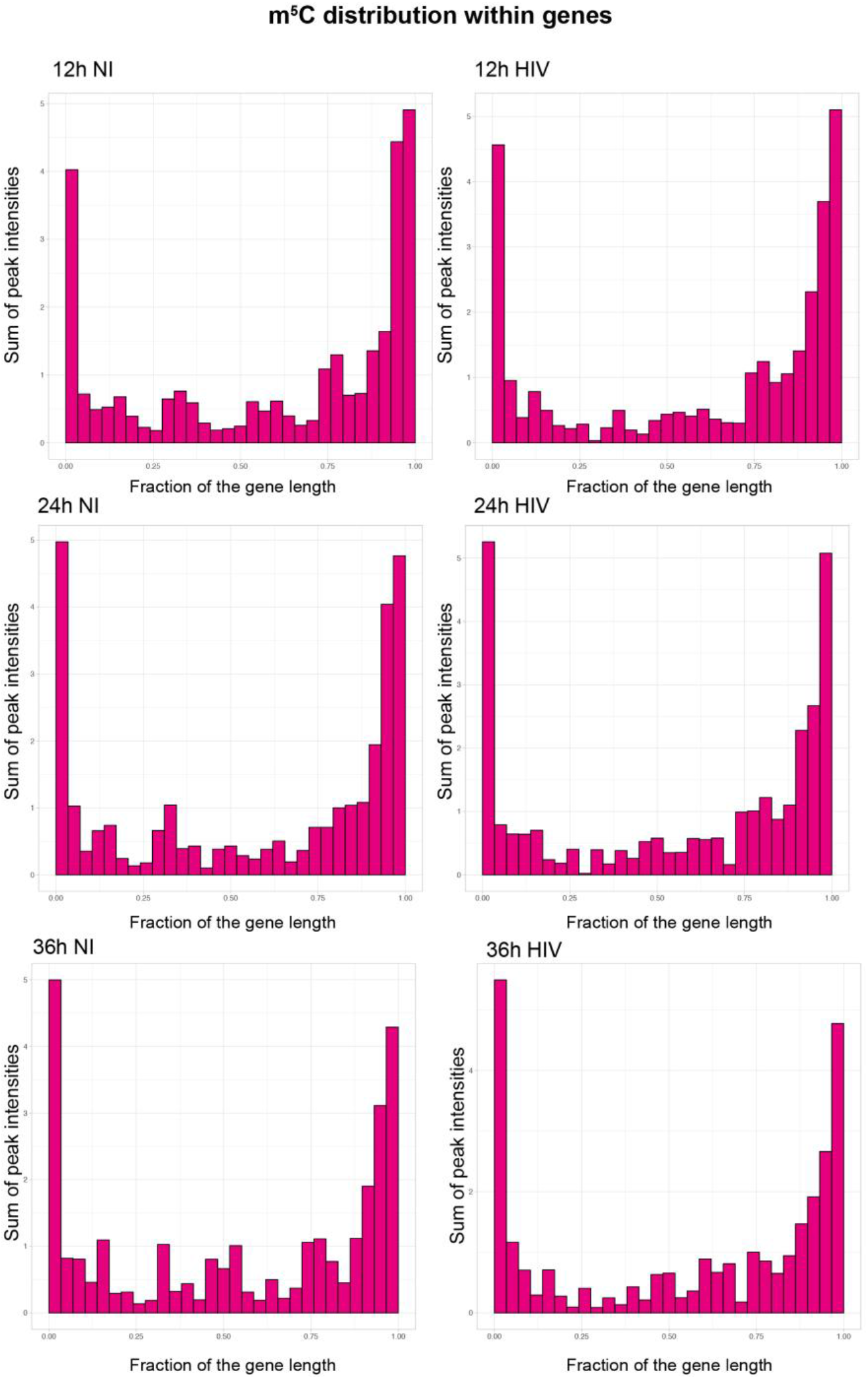
M^5^C distribution. Frequency of m^5^C methylation in each sample along genes, normalized by their gene length. Histogram plots showing on the x-axis genes normalized for their length and divided into 30 bins, and for each bin fraction of the gene, the number of m^5^C residues weighted for the methylation rate.

**Figure S7.**
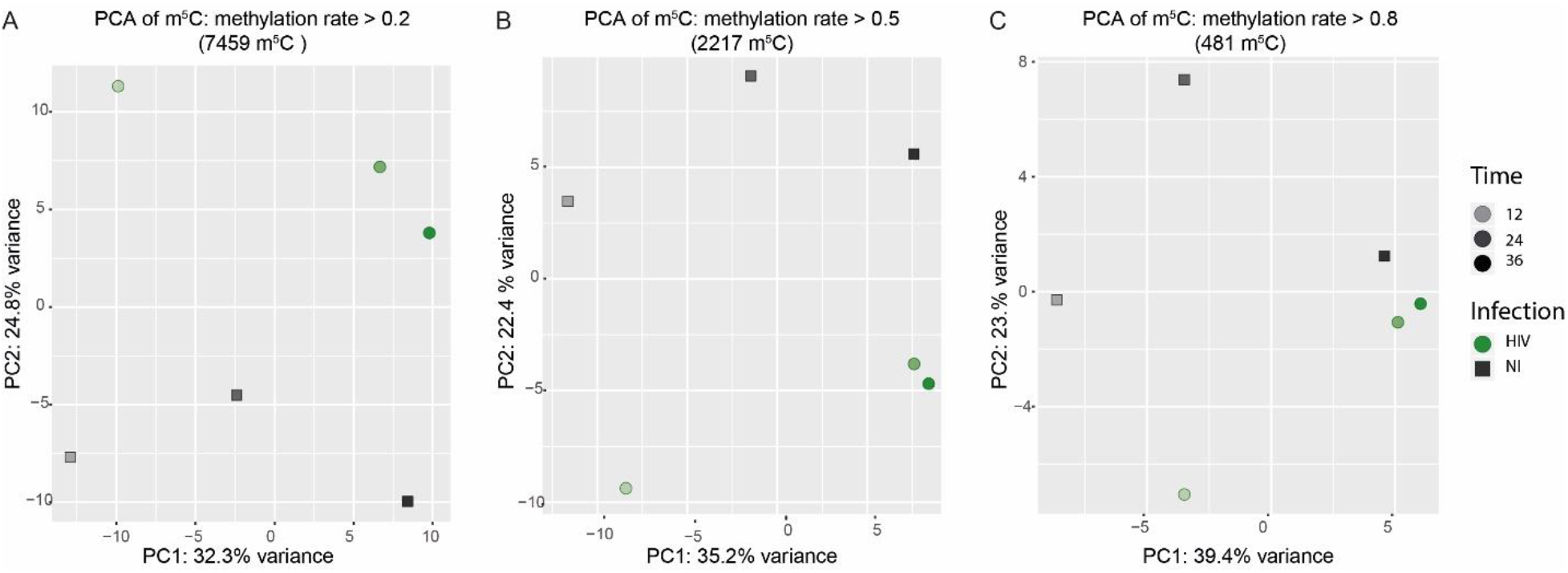
PC analysis of m^5^C across samples according to methylation rate. Different filters for methylation were considered: methylation rate >20% (A), >50% (B) and >80% (C). HIV-infected samples are represented as green filled circles, non-infected samples as grey squares. Timepoints are depicted by color shading. HIV transcripts are not included.

**Figure S8.**
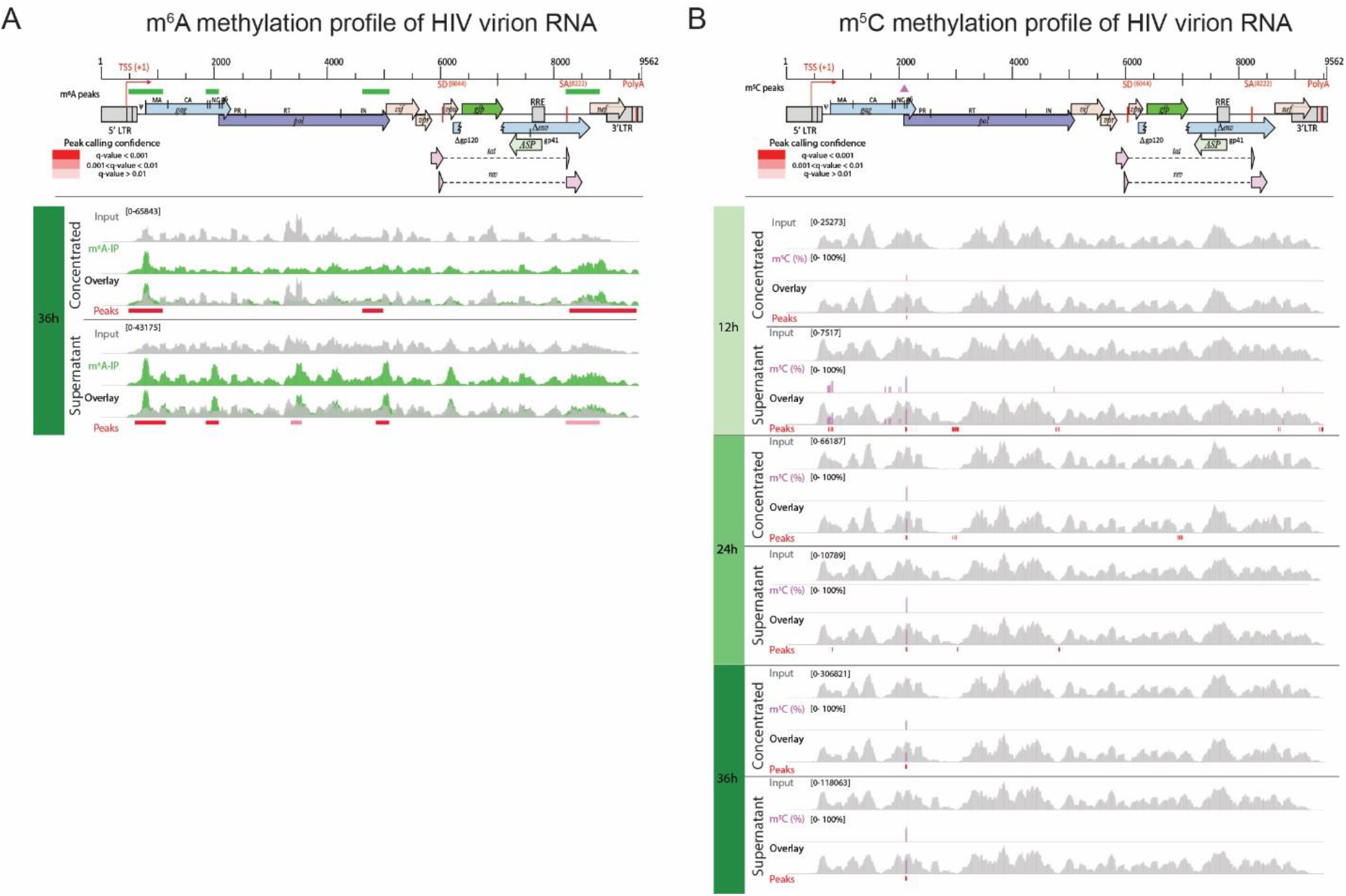
m^6^A and m^5^C profile in virions. Virion RNA was extracted either from concentrated and purified supernatant (concentrated) or from untouched supernatant (supernatant). HIV genome organization is depicted on top of the panels and common methylation marks are indicated above the genomes as horizontal green rectangles (A) or pink triangles (B), respectively. Detailed read coverage is displayed for each individual sample as tracks below the genome. (A) Identification of m^6^A peaks in HIV virion RNA at 36h post-infection. Input (gray) and m^6^A immunoprecipitated samples (green) are shown. Putative m^6^A peak calling was performed with MACS2 package after subtraction of the input background (overlay). Statistically significant peaks are highlighted by a red box, with color shading proportional to the q value (m^6^A peak track). (B) Identification of m^5^C on HIV virion RNA. Coverage of HIV genome upon conversion (gray) and detection of m^5^C (pink) are shown. M^5^C are presented as proportion of converted C. Bar height is proportional to the percentage of methylated C in the reads covering the position. The track height is set to 100%. M^5^C calling was performed with MACS2 package. Statistically significant residues are highlighted by a red box, with color shading proportional to the q value.

